# SLX4IP-mediated telomere maintenance is essential for androgen receptor-independent castration-resistant prostate cancer

**DOI:** 10.1101/2020.03.31.018846

**Authors:** Tawna L. Mangosh, Wisam N. Awadallah, Magdalena M. Grabowska, Derek J. Taylor

## Abstract

In advanced prostate cancer, resistance to androgen deprivation therapy is achieved through numerous mechanisms, including loss of the androgen receptor (AR) allowing for AR-independent growth. Therapeutic options are limited for AR-independent castration-resistant prostate cancer, and defining mechanisms critical for its survival is of utmost importance for targeting this lethal disease. Our studies have focused on defining the telomere maintenance mechanism (TMM) required for castration-resistant prostate cancer (CRPC) cell survival. TMMs are responsible for telomere elongation to instill replicative immortality and prevent senescence, with the two TMM pathways available being telomerase and alternative lengthening of telomeres (ALT). Here, we show that AR-independent CRPC exhibits ALT hallmarks and limited telomerase expression and activity, whereas AR-dependent models use telomerase for telomere maintenance. AR-independent CRPC exhibited elevated levels of SLX4IP, a protein implicated in TMM switching. SLX4IP overexpression in AR-dependent CRPC C4-2B cells promoted ALT hallmarks *in vitro*. SLX4IP knockdown in AR-independent CRPC cells (DU145 and PC-3) led to the loss of ALT hallmarks, dramatic telomere shortening, induction of senescence, and reduced tumor volume. Using an *in vitro* model of CRPC progression, induction of neuroendocrine differentiation in AR-dependent CRPC cells promoted ALT hallmarks in an SLX4IP-dependent manner. Lack of sufficient SLX4IP expression prevented ALT hallmarks rendering a TMM deficient environment, thus inducing senescence. This study demonstrates a unique reliance of AR-independent CRPC on SLX4IP-mediated ALT. Furthermore, ALT hallmark inhibition via SLX4IP induces senescence, thereby abolishing the replicative immortality of AR-independent CRPC.

## INTRODUCTION

Prostate cancer relies upon androgen receptor (AR) signaling thus androgen deprivation therapy is indicated first line and provides favorable outcomes in localized disease.^1^ However, in cases of advanced disease, prostate cancer treatment and progression is extremely complex.^1^ Androgen deprivation therapy remains the standard of care in advanced disease despite the unavoidable progression to castration-resistant prostate cancer (CRPC).^1–3^ Castration-resistance is acquired through a number of AR-dependent mechanisms such as AR mutations, amplifications, and splice variants.^4^ Resistance is also attained following loss of AR and induction of neuroendocrine differentiation (NED) allowing for AR-independent growth.^5–8^ Accordingly, second-generation antiandrogens have been developed for CRPC,^9,10^ however, they provide little benefit in CRPC patients with AR loss.^6,11^ Moreover, AR-dependent CRPC treatment with second-generation agents induces AR loss and NED.^6^ As these are complex and variable disease states, explored through *in vitro and* xenograft models, we will simply refer to these as AR-dependent and AR-independent CRPC phenotypes. Because of the lack of efficacious therapeutics and poor prognosis of AR-independent CRPC, it is crucial to identify regulatory mechanisms required for AR-independent CRPC survival.

Our studies have focused on defining telomere maintenance mechanisms (TMM) in AR-dependent and independent CRPC. Cancer cells acquire replicative immortality by activating TMM, which allows for maintained telomere length over successive cell divisions.^12–15^ Telomeres comprise the distal chromosomal ends and consist of repeating TTAGGG DNA sequences sheathed in shelterin protein complexes.^16–19^ As replicative polymerases are unable to fully replicate the ends of linear DNA,^20^ telomeric DNA of somatic cells progressively shortens with each replication cycle until a minimum threshold limit is reached provoking senescence to prevent deleterious genomic consequences.^21^ Conversely, malignancies employ TMM, through activation of telomerase or the alternative lengthening of telomeres (ALT) pathway, to evade this terminal fate and instill replicative immortality.^12–15^ Telomerase is a reverse transcriptase capable of catalyzing the addition of telomeric repeats to chromosomal ends^12^ and is the primary contributor to telomere elongation in 85-90% of cancers.^22^ ALT activation accounts for the remaining 10-15% of cancers^14^ and involves telomere-directed homologous recombination (HR) to promote elongation.^13^ The ALT mechanism remains challenging to characterize due to the inability to measure ALT activity directly, unlike the telomerase pathway.^22,23^ As such, the identification of ALT relies upon characterization of several surrogate measures including ALT-associated Promyelocytic leukemia protein (PML) bodies (APBs) found at sites of telomeric HR^24^ and extra-telomeric circular C-rich DNA byproducts of ALT called C-circles.^25^ Historically, ALT was predominantly associated with cancers with undetectable telomerase activity^14^ but emerging data indicates that telomerase and ALT can co-exist, further confounding TMM characterization.^26,27^ Despite the complicated mechanism and identification of ALT, recent evidence has begun to illuminate the underestimation of ALT’s contribution to telomere maintenance in malignancy, including prostate cancer.

Both *in vitro* and *in vivo,* telomere elongation in primary prostate cancer has been wholly attributed to telomerase.^28,29^ This is not surprising as androgens in primary disease models promote expression of the telomerase reverse transcriptase component, *TERT*.^29^ However, the primary TMM of CRPC remains unexplored. Historically, CRPC regardless of AR expression, was assumed to rely upon telomerase for telomere elongation,^28^ which is unsurprising due to the challenges surrounding ALT identification. However, a case study comparing primary and castration-resistant disease identified ALT hallmarks in a distant metastatic site, which were lacking in the primary tumor.^30^ To date, this is the only telomerase-to-ALT shift linked to disease progression, as well as the only documented occurrence of this understudied TMM in prostate cancer. Thus, understanding the acquisition of ALT in prostate cancer progression is critical for understanding the purpose of TMM shifts in this disease.

Additional examples of telomerase-to-ALT shifts have emerged in cases beyond prostate cancer progression, thereby highlighting the complexity of malignant TMM programming. For instance, cancer cells initiate telomerase-to-ALT shifts as a mechanism of resistance to telomerase inhibition.^31–34^ Despite the identification of TMM shifts, key regulators have yet to be well characterized. Recently, SLX4IP was implicated as a regulator of ALT in TMM shifts. SLX4IP interacts with SLX4, a nuclease scaffold essential for ALT-mediated telomere maintenance.^35,36^ SLX4 shuttles its associated nucleases and SLX4IP to participate in the resolution of HR junctions at the telomere following TRF2-mediated recruitment, an essential shelterin component.^35,37^ Knowledge regarding the function of SLX4IP beyond this interaction is limited but emerging data demonstrates SLX4IP differentially regulates TMM based on cancer type. Specifically, SLX4IP knockout promoted ALT in osteosarcoma models^38^ but in breast cancer models SLX4IP knockdown led to the loss of ALT hallmarks and induction of telomerase.^39^ Despite this disease-dependent difference, it is evident that SLX4IP is involved in TMM shifts though its involvement in those linked to disease progression, specifically prostate cancer, remains uninvestigated.

In order to understand the acquisition of ALT in CRPC we have characterized the primary TMM of four diverse CRPC *in vitro* models. CRPC cell lines utilizing AR-independent modes of resistance exhibit ALT hallmarks coupled with elevated SLX4IP expression, while AR-dependent models of CRPC predominantly use telomerase. Additionally, we show that overexpression of SLX4IP promotes ALT hallmarks and reduces telomerase expression and activity. In contrast, we show that SLX4IP knockdown leads to loss of ALT hallmarks in AR-independent CRPC cell lines. Consequently, these cells undergo a dramatic reduction in telomere length coupled with senescence. Despite the co-occurrence of ALT and AR-independent phenotypes, we demonstrate that the molecular programming behind each functions autonomously. Herein, we identify a unique relationship between SLX4IP and ALT that is necessary for the maintenance of telomere length and therefore replicative immortality of AR-independent CRPC.

## MATERIALS AND METHODS

### Cell Culture

Cell lines were obtained from the American Type Culture Collection and maintained at 37°C in 5% CO_2_. U2OS, HEK293T, and GP2-293 cells were cultured in DMEM (Gibco) with 10% (v/v) FBS (Sigma), 1% of Penicillin 10 000 units/mL, Streptomycin 10 000 μg/mL, and 25 μg/mL Amphotericin B mixture (Antibiotic-Antimycotic, Gibco). C4-2B, 22Rv1, DU145, and PC-3 cells were cultured in RPMI1640 (Gibco) with 10% FBS and 1% Antibiotic-Antimycotic. For androgen-depleted conditions, C4-2B cells were cultured for 12 weeks in RPMI1640, No Phenol Red (Gibco) with 10% Charcoal Stripped FBS (Sigma) and 1% Antibiotic-Antimycotic. Cell lines were subjected to MycoAlert™ PLUS Detection Kit in July 2019 (Lonza). To determine PD, cells were plated and trypsinized at 72 and 120 hours. Using trypan blue exclusion, viable cell number was determined using the Countess II (ThermoFisher).

### Stable Cell Line Generation

To generate cell lines stably overexpressing 3xFLAG-tagged-SLX4IP, a gBlock® was designed incorporating 3xFLAG tag followed by SalI cut site to the c-terminus of the SLX4IP sequence (NCBI Nucleotide) with a BamHI cut site to the n-terminus followed by bacterial codon optimization using the gBlock® Gene Fragment design tool (IDT). The gBlock® was cloned into pBABE-puro (Addgene#1764).^40^ Retrovirus was produced using Lipofectamine™ 2000 (ThermoFisher) to co-transfect pBABE-puro or pBABE-puro.SLX4IP.3xFLAG with pCMV-VSV-G (at a ratio of 6:1, Addgene#8454) into GP2-293 cells (Clontech). Virus was harvested and cleared via centrifugation at 48 hours post-transfection and infections were carried out in the presence of 10 μg/mL of polybrene (Sigma) followed by selection in 1 μg/mL (C4-2B) or 1.5 μg/mL (DU145, PC-3) of puromycin (Sigma).

To generate cell lines with stable knockdown of SLX4IP, two short hairpin RNAs (shRNA) targeting SLX4IP in the pLKO.1-puro lentiviral expression plasmid were purchased from Sigma (Clone ID NM_001009608.1-426s1c1 and NM_001009608.1-247s1c1). Lentivirus was produced using Lipofectamine™ 2000 (ThermoFisher) to co-transfect pLKO.1-puro Non-Mammalian shRNA Control (Sigma#SCH002), pLKO.1-puro.shSLX4IP.1, or pLKO.1-puro.shSLX4IP.2 with pMD2.G, pRRE, and pRSV-Rev (at a ratio of 4:1:1:1, Addgene#12259, #12251, #12253)^41^ into HEK293T cells. Viral infections and selection were carried out as described above.

### Mouse Xenograft Studies and Immunohistochemistry

*In vivo* experiments were performed with approval from the Institutional Animal Care and Use Committee at Case Western Reserve University, which is certified by the American Association of Accreditation for Laboratory Animal Care. Six-week-old male nude mice were inoculated with a 1:1 mix of Matrigel (Gibco 354234) and 1×10^6^ cells subcutaneously. Mouse weights and tumor volumes were determined three times weekly. Tumor volumes were calculated using the hemiellipsoid volume formula following Vernier caliper measurements of tumor diameters in three dimensions. At 30 days post-inoculation, tumors were excised and processed for immunohistochemistry and telomerase activity. Immunohistochemistry and hematoxylin and eosin staining was carried out as published previously.^42^ Primary antibody dilutions: SLX4IP (1:200, Sigma, HPA046372), p21 (1:500, Cell Signaling, 2947), Ki-67 (1:1 000, Sigma, HPA001164), and active caspase-3 (1:1 000, Millipore, AB3623). Three fields of view for each slide were imaged using an Olympus BX43 Upright Microscope and cells were manually counted and scored using ImageJ Software.

### RNA Isolation and Analysis

RNA was isolated using TRIzol reagent (Invitrogen), reverse transcribed using the High-Capacity RNA-to-cDNA Kit (ThermoFisher), and quantified via Quantitative real time PCR (RT-qPCR) on an Applied Biosystems™ StepOnePlus™ real time PCR system as described previously.^43^ Established primer pairs for *TERT* and for *GAPDH* were used.^43^

### Western Blotting Analysis

Western blotting analysis was completed as previously described.^43^ Briefly, 50 μg of lysate was resolved on a 4-20% Mini-PROTEAN® TGX™ Gel (BioRad) and transferred to Nitrocellulose membrane (Millipore) using the Trans-Blot® Turbo™ (BioRad). Blots were blocked in 5% milk in TBST and incubated overnight at 4°C with primary antibody. Blots were incubated for one hour at room temperature with either HRP-conjugated (Santa Cruz), IRDye 800CW or IRDye 680RD (LI-COR) secondary antibodies diluted to 1:10 000. Development with ProtoGlow ECL reagent (National Diagnostics) was used when necessary. Blots were imaged on an Odyssey® Fc Imaging System (LI-COR) and quantified using Image Studio™ 5.2 Software. Primary antibody dilutions: 1:20 000 GAPDH (Cell Signaling, 2118), 1:5 000 SLX4IP (Sigma, HPA046372), 1:5 000 FLAG (Sigma, F1804), 1:10 000 AR (Santa Cruz, sc-816), 1:5 000 FOXA1 (Abcam, ab23738), 1:3 000 ENO2 (Cell Signaling, 9536), 1:5 000 p21 (Cell Signaling, 2947).

### Telomere Repeat Amplification Protocol

The real-time Q-TRAP assay was performed as previously published.^44^ Tumor lysates were prepared following homogenization with Kontes® PELLET PESTLE® Grinders (Kimble) under similar lysis conditions. Reactions were performed on an Applied Biosystems™ StepOnePlus™ real time PCR system. Collected Ct values were then converted to relative telomerase activity (RTA) units using the telomerase-positive standard curve generated as previously described.

### Immunofluorescence-Fluorescence *in situ* Hybridization Analysis

A previously published immunofluorescence-fluorescence *in situ* hybridization protocol was carried out on cells seeded on sterile glass coverslips and tumor sections.^43^ Coverslips were incubated with PML antibody (Santa Cruz, sc-5621) diluted 1:200 for one hour at room temperature. Alexa-488® conjugated anti-rabbit secondary antibody (Jackson) was diluted 1:100 and added to coverslips for overnight incubation at 4°C. Hybridization with 333ng/mL of a TelC-Cy5-labeled peptide nucleic acid (PNA) oligonucleotide telomere probe (N-CCTAACCTAACCTAA-C, PNA BIO) in PNA buffer (10mM Tris pH 7.5, 70% formamide) was carried out at 95°C for 5 min followed by overnight incubation at room temperature. Coverslips were stained with 2 μg/mL of DAPI for 10 min and mounted with Fluoromount-G (ThermoFisher). Stained slides were imaged using the Leica HyVolution SP8 gated STED Microscope and manually scored for co-localization events and cell count using LasX Software.

### C-circle Isolation and Analysis

A previously published C-circle dot blot assay was performed with minor modifications.^45^ 200 ng of DNA was incubated under conditions described in the previously published protocol. Reactions were blotted onto Hybond® membranes (ThermoFisher) and UV cross-linked (Spectrolinker™ XL-1000, Spectronics). Membranes were hybridized with 30μg/mL of Digoxigenin (DIG) conjugated telomere probe (CCCTAACCCTAACCCTAACCCTAA-DIG, IDT) in DIG Easy Hyb buffer (Roche) overnight at 42°C. 2x saline-sodium citrate (SSC) buffer with 0.1% SDS was used to remove excess probe. Membranes were blocked in DIG Blocking Buffer (Roche) for 30 min followed by incubation with 1:10 000 dilution of Digoxigenin-AP, Fab fragments (Roche) for 3 hours at room temperature. DIG Wash buffer (Roche) was used to remove excess antibody and amplified C-circles were detected with CDP-Star® kit (Roche). Membranes were imaged with the Odyssey® Fc Imager (LI-COR) and signal intensity ratios calculated as previously described using Image Studio 5.2 Software.

### Telomere Restriction Fragment Analysis

Genomic DNA was harvested using the Genelute™ Mammalian Genomic DNA Miniprep Kit (Sigma) according to manufacturer’s instructions and telomere restriction fragment analysis was carried out as previously published.^43^ Membranes were hybridized, prepared, and imaged as described in the above C-circle analysis protocol.

### Senescence-associated β-galactosidase Activity Staining

Cells were seeded on sterile glass coverslips and grown to 80% confluency. Senescence β-galactosidase staining kit was used following the manufacturer’s protocol (Cell Signaling Technologies). Slides were imaged using an Olympus BX43 Upright Microscope and cells were manually counted and scored using ImageJ Software.

### Statistical Analysis

Using GraphPad Prism 8, statistics were performed using two-tailed Student’s *t* test (*in vitro* and *in vivo* data comparing two groups) or one-way ANOVA with multiple comparisons (*in vitro* data comparing at least three groups). Patient expression data was analyzed through the PCTA based on disease course using Ranksums-test between mCRPC and primary subsets (www.thepcta.org). *p* values less than 0.05 were considered statistically significant. All *in vitro* data are represented as mean values from three independent experiments performed in triplicate unless otherwise noted in respective figure legends.

## RESULTS

### AR-independent CRPC exhibits ALT hallmarks and limited telomerase activity

Due to the underestimation of ALT’s contribution to telomere maintenance, we sought to determine if primary TMM correlated with AR expression. Four *in vitro* CRPC models were interrogated for telomerase expression, activity, and ALT hallmarks. C4-2B and 22Rv1 cell lines retain AR expression^46,47^ while DU145 and PC-3 cells exhibit AR loss.^48,49^ Additionally, PC-3 and DU145 cells exhibit markers of NED, a phenotype commonly observed with CRPC disease progression following androgen deprivation.^50–52^ Relative mRNA expression of *TERT and* telomerase activity across cell lines was analyzed in tandem with ALT-positive U2OS control (Fig. 1A and B). Interestingly, only AR-independent DU145 and PC-3 cells demonstrated both relatively low *TERT* expression and telomerase activity.

**Figure 1:**
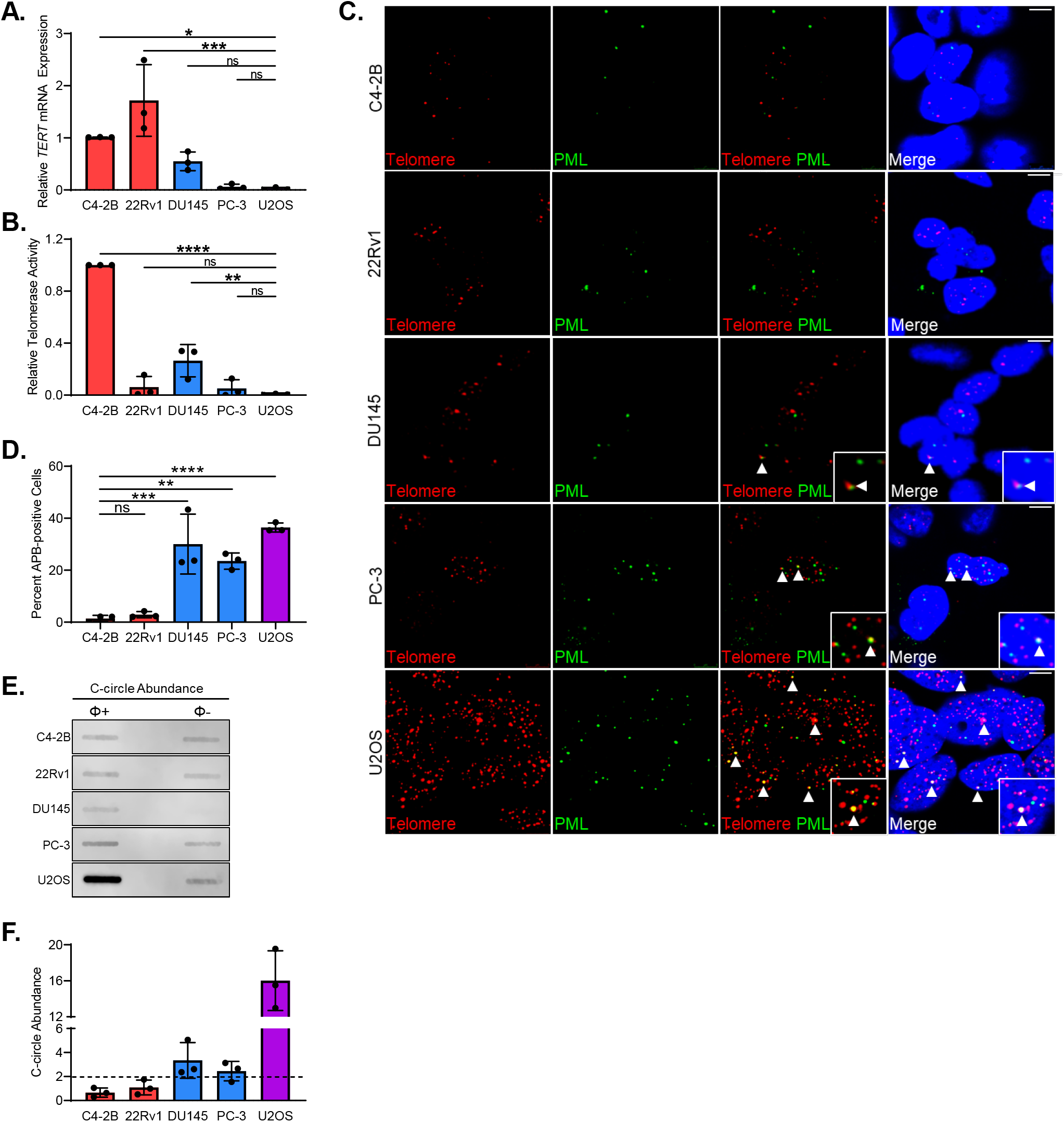
AR-independent CRPC cell lines exhibit ALT hallmarks. **(A)** Relative *TERT* mRNA expression and **(B)** relative telomerase activity across four CRPC cell lines, C4-2B and 22Rv1 which retain AR expression and DU145 and PC-3 which exhibit AR loss. ALT-positive U2OS cells were included as control. **(C)** Representative IF-FISH images demonstrating the presence or absence of APBs at telomeres (arrowheads) in CRPC cell lines and U2OS control. Scale bar: 5 μm. **(D)** Quantification of IF-FISH images for percent APB-positive cells defined as cells with at least one or more telomeric foci (red) localizing with PML protein (green). **(E)** Representative dot blot demonstrating the presence of telomeric C-circles in each CRPC cell line. Φ+ lane indicates reactions incubated with Φ29 polymerase for telomeric C-circle amplification. Φ-lane indicates control reaction lacking polymerase to account for background telomeric signal. **(F)** Quantification of C-circle abundance defined as the signal ratio of Φ+ reaction to Φ-reaction. Dotted line at y=2 indicates threshold signal ratio suggesting ALT activity. Data represented smean±SD; n=3; *p<0.05; **p<0.01; ***p<0.001; ****p<0.0001.

Based on this result, we reasoned that ALT may be engaged to compensate for the minimal telomerase contribution to telomere maintenance in this AR-independent environment. As ALT activity cannot be directly measured, we evaluated the abundance of APBs and C-circles.^24,25^ First, APBs were visualized via immunofluorescence-fluorescence *in situ* hybridization for ALT-associated PML protein localizing at the telomere, which is highly indicative of ALT-mediated HR events.^24^ DU145 and PC-3 cell lines had a similar proportion of APB-positive cells as the ALT-positive control; however, the C4-2B and 22Rv1 cells retaining AR had significantly fewer APB-positive cells (Fig. 1C and D). Second, telomeric C-circles that are byproducts of ALT-related events were evaluated via C-circle Amplification (CCA).^45^ A signal ratio greater than or equal to two when comparing the C-circle amplification reaction to control is highly indicative of ALT activity^45^ and only cell lines with AR loss reached this threshold (Fig. 1E and F). Taken together these data indicate CRPC cell lines with AR loss exhibit ALT hallmarks with limited telomerase expression and activity. In contrast, CRPC cell lines retaining AR expression do not exhibit the ALT phenotype supporting telomerase as the major TMM player.

### Elevated SLX4IP expression correlates with AR loss and ALT Hallmarks in CRPC

The CRPC case study demonstrating ALT hallmarks exhibited loss of chromatin remodeler ATRX.^30^ While ATRX deletions are potential ALT initiating events, these genomic alterations are not universally causative of ALT across *in vitro* models.^53,54^ Mutational analysis of ATRX^55,56^ and mRNA and protein expression across CRPC cell lines revealed DU145 and PC-3 cells, which display ALT hallmarks, retain wild-type ATRX expression (Supplemental Fig. 1A and B). While ATRX loss may be responsible for ALT initiation in some cases, it is noncontributory in our CRPC *in vitro* models. Therefore, additional ALT regulators were investigated.

The relatively uncharacterized protein, SLX4IP, was identified as a promoter of TMM shifts and ALT-mediated telomere maintenance in breast cancer models; however, its role in CRPC is unknown.^39^ Analysis of relative SLX4IP protein expression revealed DU145 and PC-3 cells, which exhibit ALT hallmarks and AR loss, had significantly higher SLX4IP expression compared to their AR-dependent counterparts lacking ALT hallmarks (Fig. 2A and B). Consistently, inspection of SLX4IP expression data from the Prostate Cancer Transcriptome Atlas^57^ revealed significantly higher expression in metastatic CRPC (mCRPC) patients when compared to those with primary disease (Fig. 2C). The correlation between elevated SLX4IP expression and ALT hallmarks and known interactions with ALT-associated SLX4 and TRF-2 supports SLX4IP as a potential regulator of TMM in CRPC.^35,37^

**Figure 2:**
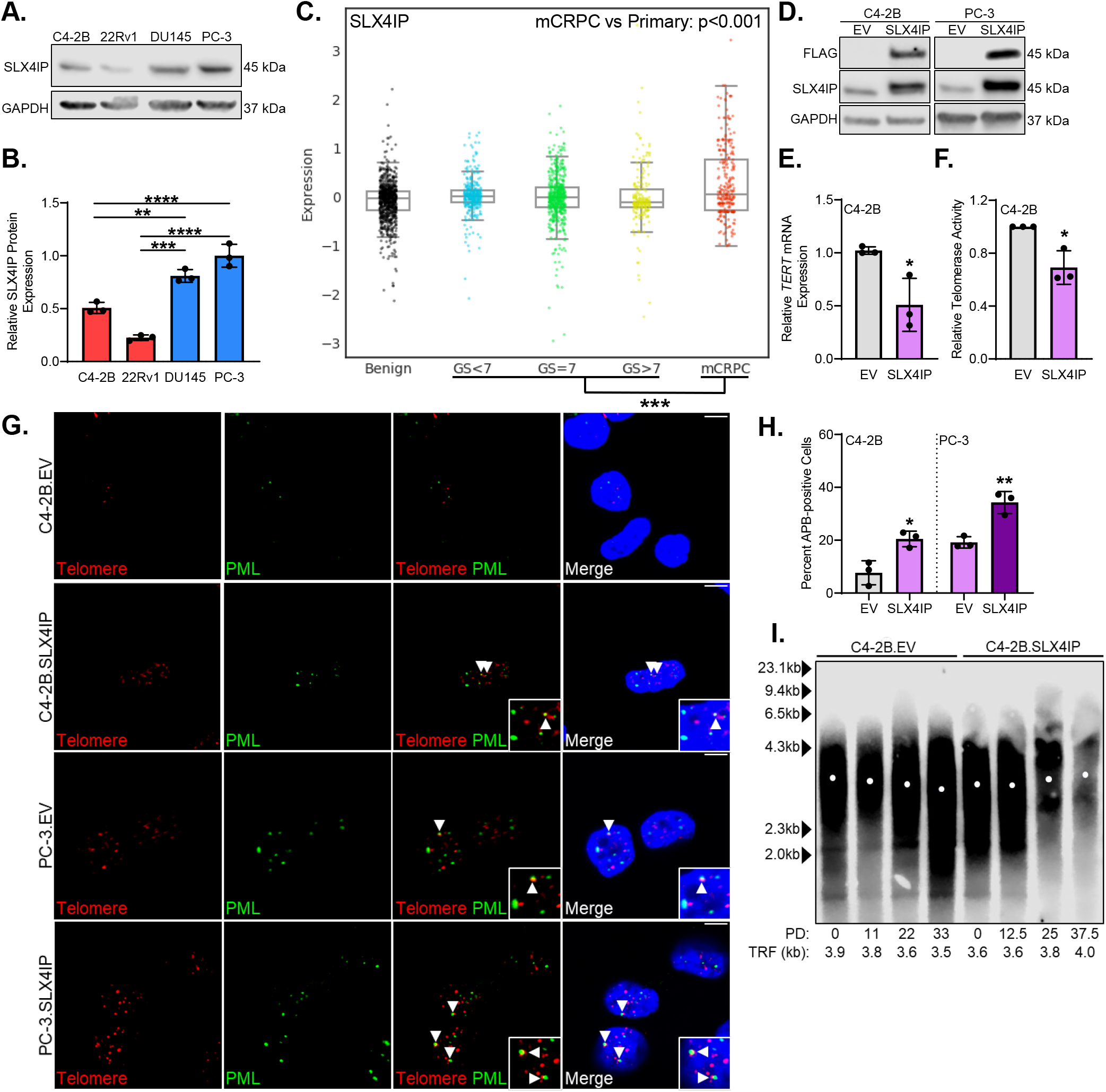
Elevated SLX4IP expression correlates with ALT hallmarks in CRPC. **(A)** Relative SLX4IP protein expression across CRPC cell lines. **(B)** Quantification of (A). **(C)** SLX4IP expression data from the PCTA arranged based upon patient diagnosis of benign disease, primary disease and Gleason score, or mCRPC. Ranksums-test was used to compare primary (GS<7,=7,>7) versus mCRPC cohorts. **(D)** Confirmation of stable overexpression of SLX4IP in C4-2B and PC-3 cells. EV: Empty Vector. **(E)** Relative *TERT* mRNA expression and **(F)** relative telomerase activity following stable SLX4IP overexpression in C4-2B cells. **(G)** Representative IF-FISH images demonstrating the presence or absence of APBs at telomeres (arrowheads) in C4-2B and PC-3 cells stably overexpressing SLX4IP. Scale bar: 5 μm. **(H)** Quantification of IF-FISH images for percent APB-positive cells in SLX4IP overexpressing C4-2B and PC-3 cells compared to control. **(I)** Telomere restriction fragment analysis demonstrating telomere length changes over 45 days in C4-2B cells overexpressing SLX4IP versus control. PD: Calculated population doublings. Data represented as mean+SD; n=3; *p<0.05; **p<0.01; ***p<0.001; ****p<0.0001.

Therefore, the ability of SLX4IP to promote ALT hallmarks was investigated in CRPC cell lines. A 3xFLAG-tagged SLX4IP construct was stably overexpressed via retroviral transduction in C4-2B (C4-2B.SLX4IP) and PC-3 cells (PC-3.SLX4IP) (Fig. 2D). In C4-2B.SLX4IP cells, a significant reduction in *TERT* expression and telomerase activity was observed (Fig. 2E and F). Unsurprisingly, this effect was diminished in PC-3 cells that have limited telomerase expression and activity at baseline (Supplemental Fig. 2A and B). As predicted, SLX4IP overexpression coincided with a significant increase in the percent of APB-positive cells in both C4-2B.SLX4IP and PC-3.SLX4IP cell populations (Fig. 2G and H). Lastly, SLX4IP overexpression resulted in a significant elevation in C-circle signal ratio promoting C4-2B.SLX4IP cells to beyond the threshold that is indicative of ALT activity (Supplemental Fig. 2C and D).^45^ Though a modest elevation in C-circle abundance was noted in PC-3.SLX4IP cells as well, the relative change was insignificant (Supplemental Fig. 2C and D). Together, these data indicate SLX4IP overexpression is capable of promoting ALT hallmarks.

These data support the role of SLX4IP in inducing ALT hallmarks but it is unclear how this translates to telomere length changes. After defining the population doubling (PD) time (Supplemental Fig. 2E and F), telomere restriction fragment (TRF) analysis revealed control C4-2B and PC-3 cells exhibited modest telomere length shortening over 45 days (Fig. 2I and Supplemental Fig. 2G). However, this telomere shortening was abrogated with SLX4IP overexpression, further suggesting telomeres engaged in additional elongation events, potentially due to the promotion of ALT hallmarks via SLX4IP overexpression.

Here we establish elevated SLX4IP enhances ALT hallmarks with telomerase suppression in C4-2B cells and exacerbates ALT hallmarks from baseline in PC-3 cells. Moreover, these phenotypic changes are coupled to telomeric length preservation, suggesting SLX4IP is sufficient for promoting functional telomeric elongation events in CRPC.

### SLX4IP is essential for ALT hallmarks in AR-independent CRPC

To identify the effects of SLX4IP loss on telomere maintenance in AR-independent CRPC, stable cell lines with SLX4IP knockdown (KD.1, KD.2) and non-targeting control (NS) derived from PC-3 (Fig. 3A and B) and DU145 cells were generated using lentiviral transduction. SLX4IP knockdown in PC-3 cells led to a dramatic reduction in percent of APB-positive cells (Fig. 3C and D). Furthermore, SLX4IP knockdown was sufficient to significantly reduce C-circle abundance in PC-3 cells (Supplemental Fig. 3A and B). With the loss of ALT hallmarks, we reasoned SLX4IP knockdown may trigger a compensatory induction of telomerase to address limited ALT involvement in telomere length maintenance.^58^ However, *TERT* expression and telomerase activity was not increased in PC-3.KD.1 and KD.2 cells (Fig. 3E and F). These data suggest reductions in SLX4IP expression are sufficient to render a TMM deficient telomeric environment in PC-3 cells.

**Figure 3:**
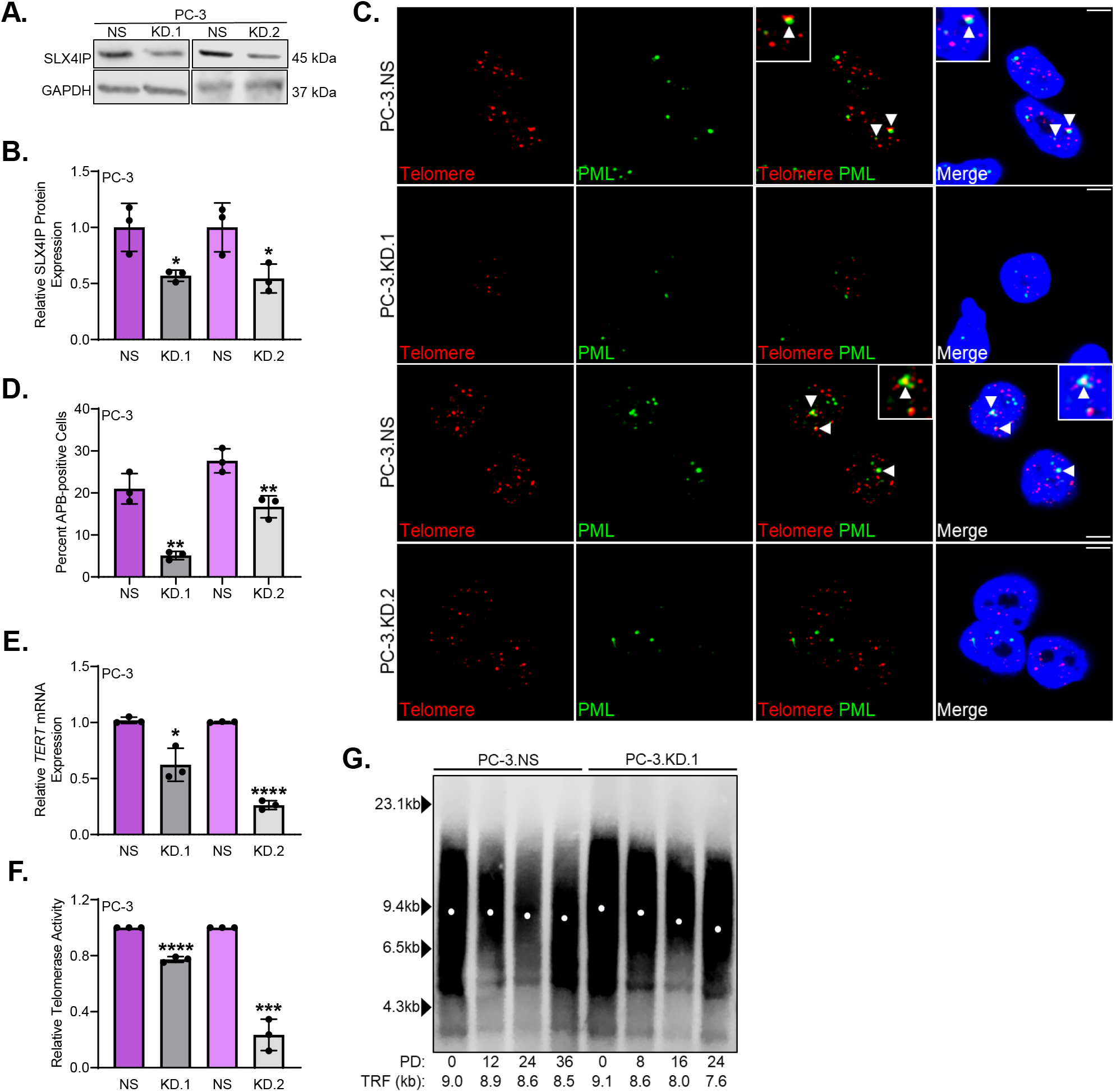
SLX4IP knockdown is accompanied by ALT hallmark disappearance and telomere shortening. **(A)** Confirmation of stable SLX4IP knockdown using two shRNAs (KD.1, KD.2) in PC-3 cells with scrambled shRNA control (NS) at the protein level. **(B)** Quantification of (A). **(C)** Representative IF-FISH images demonstrating the presence or absence of APBs at telomeres (arrowheads) in PC-3 cells with SLX4IP knockdown. Scale bar: 5 μm. **(D)** Quantification of IF-FISH images for percent of APB-positive cells. **(E)** Relative *TERT* mRNA expression and **(F)** relative telomerase activity following SLX4IP knockdown in PC-3 cells. **(G)** Telomere restriction fragment analysis demonstrating telomere length changes over 45 days in PC-3 cells with SLX4IP knockdown (KD.1) versus control. PD: Calculated population doublings. Data represented as mean+SD; n=3; *p<0.05; **p<0.01; ***p<0.001; ****p<0.0001.

Cells with insufficient TMM will exhibit more rapid telomere shortening than their parental counterpart with an intact TMM.^34^ To test this, telomere length changes via TRF were evaluated over 45 days coupled to PD time determination. Calculation of PD time revealed a dramatic reduction in cell number overtime in PC-3.KD.1 and KD.2 cells suggesting an impairment in proliferative capacity, a phenotype observed with TMM inhibition or loss (Supplemental Fig. 3C and D).^21,59^ In addition, PC-3.KD.1 and KD.2 cells demonstrated an average telomere length shortening of ∼1 500 to 2 400 bp over 24 PDs, whereas control cells exhibited a more modest shortening of only ∼400 to 500 bp in the equivalent number of PD (Fig. 3G and Supplemental Fig. 3E). Cells unable to sufficiently maintain telomeres are programmed to senesce to prevent loss of genomic information once a minimum threshold length is reached.^21,59^ Because PC-3.KD.1 and KD.2 cells demonstrate a telomeric phenotype with insufficient TMM, substantial telomere shortening, and limited proliferative capacity it is likely that senescence would be invoked. As expected, senescence-associated β-galactosidase staining^60^ revealed a considerable increase in the percent of β-galactosidase-positively (β-gal-positive) stained cells (Fig. 4A and B). Elevations in relative p21 protein expression, a senescence-associated marker^61^, further corroborated this phenotype (Fig. 4C and D).

**Figure 4:**
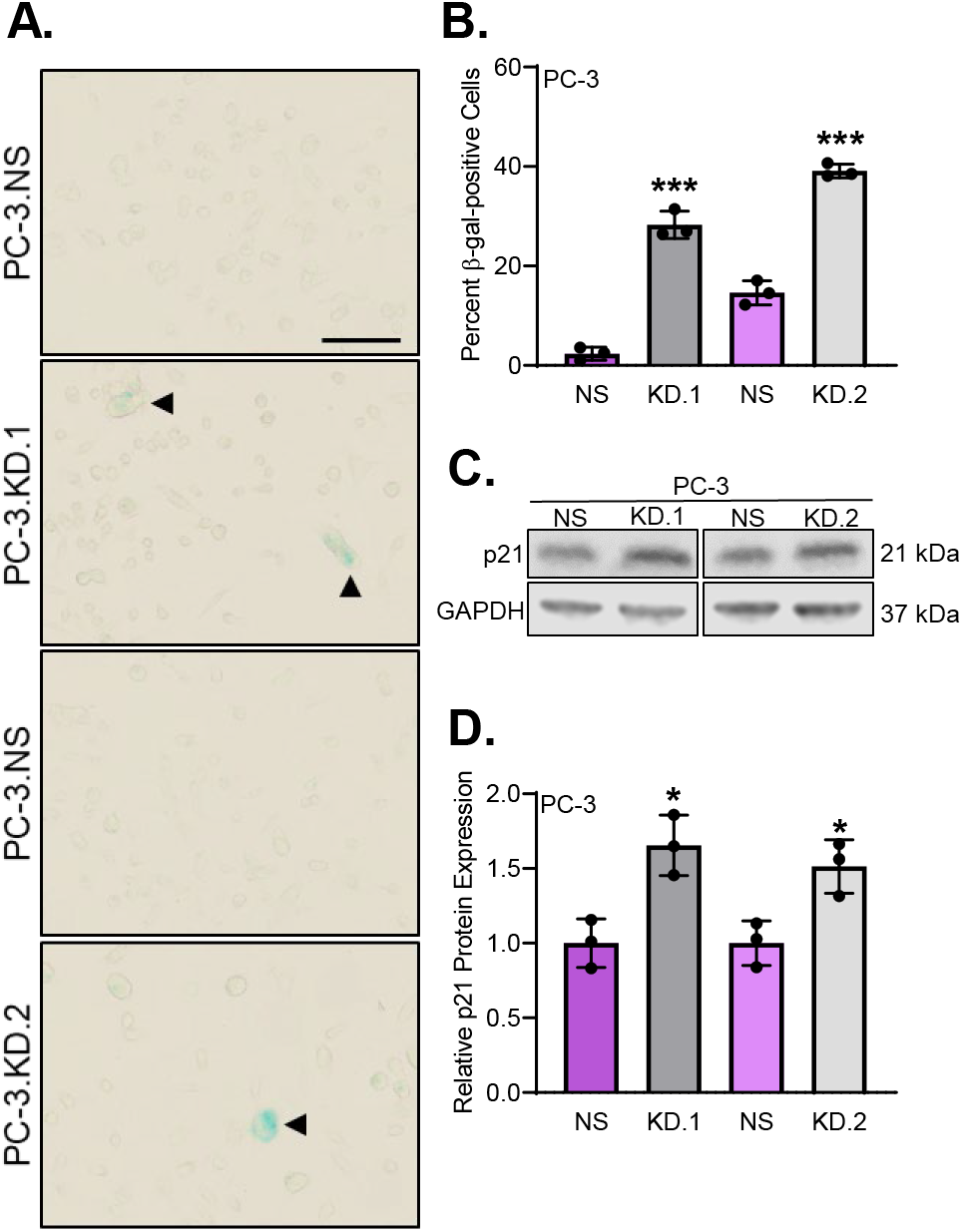
Telomere shortening following SLX4IP knockdown induces senescence. **(A)** Representative bright field images demonstrating β-galactosidase staining (arrowheads) for senescence in PC-3 cells with stable knockdown of SLX4IP. Scale bar: 20 μm. **(B)** Quantification of (A). **(C)** Relative p21 expression following knockdown of SLX4IP in PC-3 cells. **(D)** Quantification of (C). Data represented as mean+SD; n=3; *p<0.05; **p<0.01; ***p<0.001; ****p<0.0001.

In contrast to PC-3 cells, SLX4IP knockdown in DU145 cells (Supplemental Fig. 4A and B) revealed a compensatory increase in telomerase expression and activity coupled with a reduction in APB-positive cells (Supplemental Fig. 4C, D, E, and F). Though these cells have the ability to induce telomerase expression and promote functional assembly of the enzyme, a significant impairment in PD time and modest telomere shortening compared to control was observed (Supplemental Fig. 5A and B). Moreover, SLX4IP knockdown in DU145 cells led to a prominent increase in both β-gal-positively stained cells and relative p21 protein expression, demonstrating the induction of senescence despite the compensatory induction of telomerase observed (Supplemental Fig. 5C, D, E, and F).

Loss of ALT hallmarks via SLX4IP knockdown revealed a senescent phenotype in both PC-3 and DU145 cells lacking AR expression. Remarkably, the compensatory induction of telomerase observed in only DU145 cells was unable to prevent telomere shortening and senescence. These data support ALT hallmarks are necessary for adequate telomere maintenance in AR-independent CRPC. Additionally, these data identify SLX4IP is an essential regulator of ALT hallmarks in this AR-independent environment.

To ascertain if loss of ALT hallmarks via SLX4IP knockdown is maintained *in vivo,* male nude mice were inoculated subcutaneously with PC-3.NS or PC-3.KD.1 cells and tumor volumes subsequently monitored over 30 days. Immunohistochemistry revealed that SLX4IP knockdown led to a significant increase in the percent of p21-positively stained cells (Fig. 5A and B). Consequently, this effect translated to a significant reduction in average tumor volume compared to control at time of sacrifice (Fig. 5C). As no considerable difference in Ki-67 or cleaved caspase-3 staining between groups was observed the senescent phenotype is the most likely contributor to the observed tumor volume reductions (Fig. 5A). Moreover, SLX4IP knockdown resulted in the disappearance of APB-positive cells (Fig. 5D and E). Like our *in vitro* studies, evaluation of telomerase activity among xenografts did not reveal a compensatory increase in telomerase activity with SLX4IP knockdown (Fig. 5F). Together, these *in vivo* results solidify SLX4IP as a necessary mediator of ALT hallmarks in AR-independent CRPC. Without sufficient SLX4IP, senescence is induced *in vivo* translating to a significant reduction in tumor burden.

**Figure 5:**
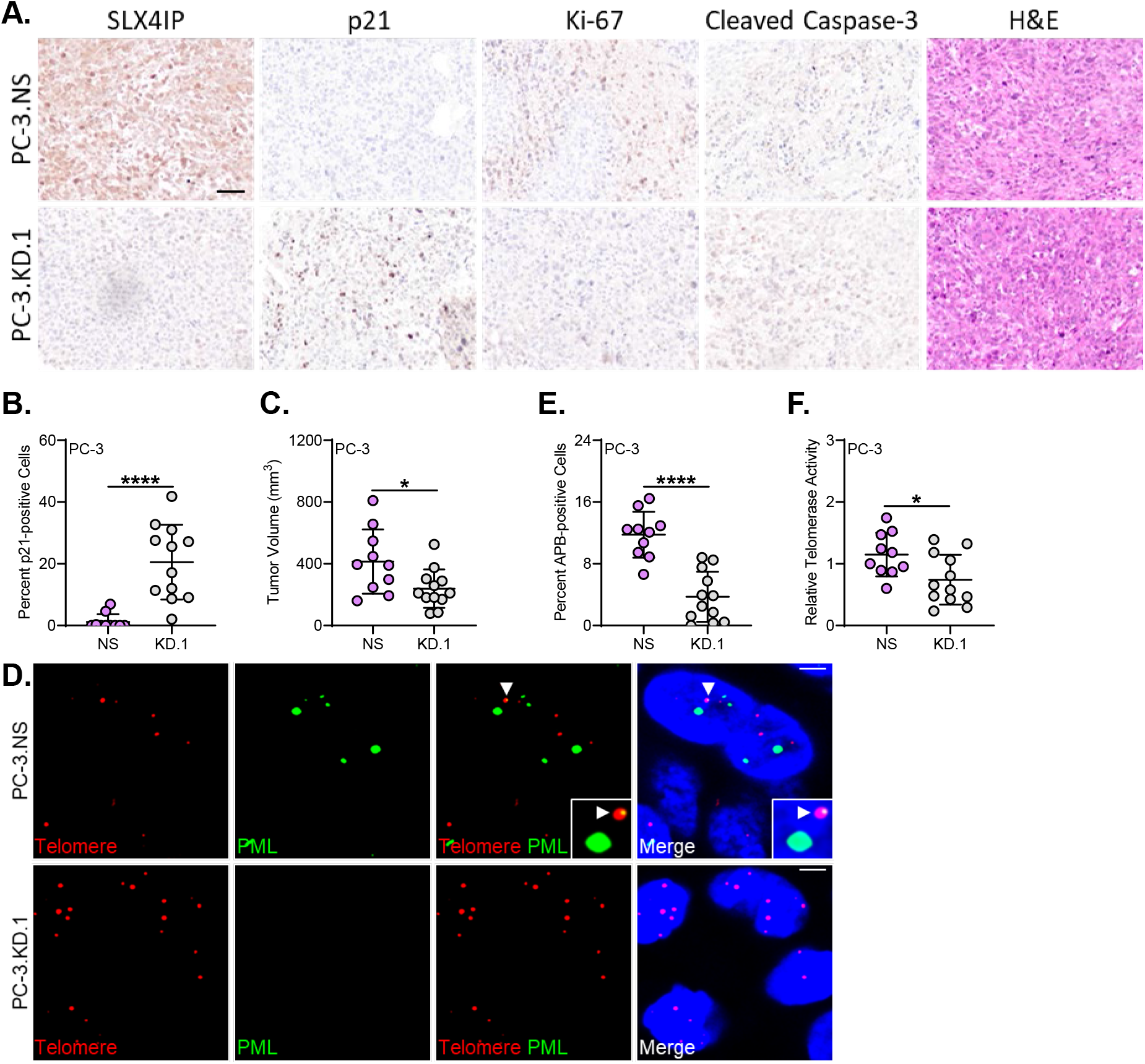
SLX4IP knockdown in AR-independent CRPC leads to reduced tumor volume, ALT hallmark loss, and senescence *in vivo*. **(A)** Representative immunohistochemistry 40X images demonstrating SLX4IP, p21, Ki-67, cleaved caspase-3, and H&E staining from PC-3.NS and KD.1 xenograft sections. Scale bar: 60 μm. **(B)** Quantification of p21 staining in (A) for percent p21-positive cells. **(C)** Average tumor volume of PC-3.NS (n=12) and KD.1 (n=10) on day 30. **(D)** Representative IF-FISH images demonstrating the presence or absence of APBs at telomeres (arrowheads) in PC-3 xenografts with SLX4IP knockdown. Scale bar: 5 μm. **(E)** Quantification of IF-FISH images for percent APB-positive cells. **(F)** Relative telomerase activity following SLX4IP knockdown in PC-3 xenografts. Data represented as mean+SD; *p<0.05; **p<0.01; ***p<0.001; ****p<0.0001.

### Androgen deprivation promotes ALT hallmarks in an SLX4IP-dependent manner

Androgen deprivation therapy triggers therapy-acquired NED and AR loss in patients.^3,50,62,63^ Similarly, AR-dependent cell lines undergo NED when placed in growth media lacking androgens.^64,65,66^ Because ALT hallmarks were identified in a CRPC patient only after androgen deprivation therapy and we identified a unique relationship between ALT and AR-independent CRPC, we reasoned AR-dependent CRPC cells lacking ALT hallmarks grown in androgen-deprived conditions may lose telomerase expression and activity and gain ALT hallmarks during NED. Moreover, this promotion of ALT hallmarks is expected to be reliant on SLX4IP. To address this question, C4-2B cells with stable knockdown of SLX4IP (KD.1, KD.2) and non-targeting control (NS) were generated and placed in charcoal-stripped media to mimic androgen deprivation and induce NED.^66^

Analysis of SLX4IP expression revealed a modest yet significant increase in SLX4IP expression when comparing C4-2B.NS cells cultured in charcoal-stripped media (+CSS) for 12 weeks, hypothesized to engage ALT, to their normal growth media (fetal bovine serum, +FBS) counterparts (Fig. 6A and B). ENO2 expression, a marker of NED, across groups confirmed the previously published transition was initiated under androgen-depleted conditions (Fig. 6A and C).^67^ Notably, a difference in ENO2 expression was not observed between cell lines grown in charcoal-stripped media indicating SLX4IP itself is not responsible for inducing NED.

**Figure 6:**
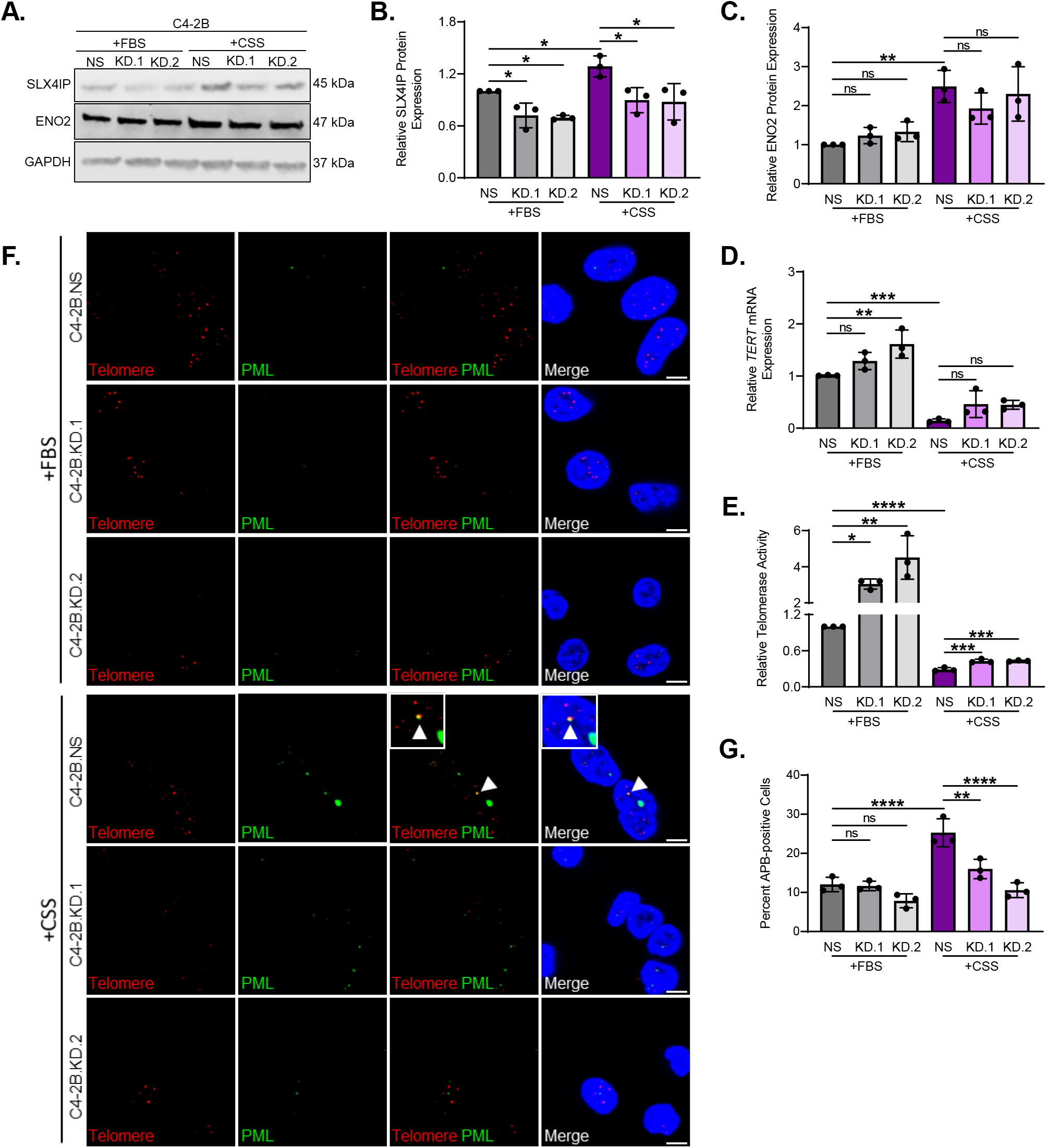
Androgen deprivation triggers SLX4IP-dependent ALT hallmarks. **(A)** Confirmation of stable SLX4IP knockdown using two shRNAs (KD.1, KD.2) in C4-2B cells with scrambled shRNA control (NS) and confirmation of NED via ENO2 expression following growth in androgen deprived conditions (+CSS) versus normal growth media (+FBS). **(B)** Quantification of SLX4IP and **(C)** ENO2 expression in (A). **(D)** Relative *TERT* mRNA expression and **(C)** relative telomerase activity following SLX4IP knockdown in C4-2B cells grown in +FBS or +CSS conditions. **(E)** Representative IF-FISH images demonstrating the presence or absence of APBs at telomeres (arrowheads) in C4-2B cells with SLX4IP knockdown. Scale bar: 5 μm. **(F)** Quantification of IF-FISH images for percent APB-positive cells. Data represented as mean+SD; n=3; *p<0.05; **p<0.01; ***p<0.001; ****p<0.0001.

Next, *TERT* expression and telomerase activity were evaluated and as expected, relatively high telomerase expression and activity was noted in complete media; however, with charcoal stripped media, all cell lines exhibited a significant reduction in both (Fig. 6D and E). As expected, few APB-positive cells were identified in C4-2B.NS, KD.1, and KD.2 cells grown in complete media but following charcoal-stripped media growth, and resulting NED, C4-2B.NS cells exhibited a significant increase in the percent of APB-positive cells (Fig. 6F and G). This increase in APBs was abrogated with SLX4IP knockdown (Fig. 6F and G). Together these data support C4-2B cells shift from high telomerase expression and activity to low telomerase expression and activity with ALT hallmarks under androgen deprivation. Without sufficient SLX4IP expression, this TMM shift cannot occur.

With SLX4IP knockdown, a reduction of ALT hallmarks combined with low telomerase expression and activity makes senescence an expected fate. Indeed, β-galactosidase staining revealed a significant increase in β-gal-positive cells with SLX4IP knockdown compared to C4-2B.NS cells grown in charcoal stripped serum (Fig. 7A and B). Additionally, p21 protein expression was elevated in SLX4IP knockdown cells cultured in charcoal stripped serum media (Fig. 7C and D). Therefore, we propose with androgen deprivation, AR-dependent CRPC can lose telomerase expression and activity and gain ALT hallmarks in an SLX4IP-dependent manner. Without sufficient SLX4IP expression and promotion of ALT hallmarks, senescence is induced in this *in vitro* model of disease progression following androgen deprivation.

**Figure 7:**
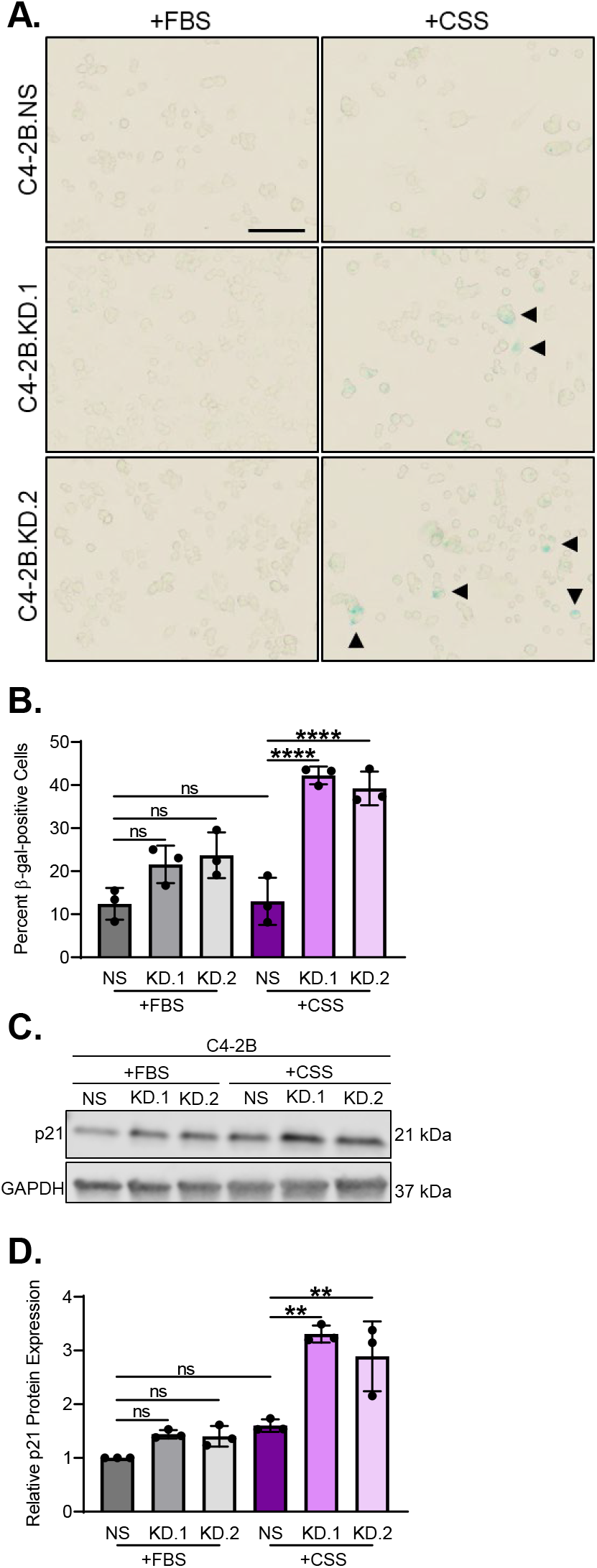
Lack of ALT hallmarks triggers senescence following androgen deprivation. **(A)** Representative bright field images demonstrating β-galactosidase staining (arrowheads) for senescence in C4-2B cells with stable knockdown of SLX4IP. Scale bar: 20 μm. **(B)** Quantification of (A). **(C)** Relative p21 expression following knockdown of SLX4IP in C4-2B cells grown in +CSS and +FBS. **(D)** Quantification of (C). Data represented as mean+SD; n=3; *p<0.05; **p<0.01; ***p<0.001; ****p<0.0001.

### ALT inhibition via SLX4IP does not prevent AR-independent CRPC expression changes

As we observed that SLX4IP is required for NED-associated ALT hallmarks, we next tested if SLX4IP overexpression is sufficient to induce NED. We again used C4-2B cells, which can undergo NED in response to androgen deprivation.^66^ Corroborating our previous result, analysis of protein expression changes in C4-2B cells with transient SLX4IP overexpression did not demonstrate a significant reduction in AR and FOXA1, a pioneering transcription factor of AR known to suppress NED^68^, or induction of ENO2 expression (Supplemental Fig. 6A and B). Moreover, SLX4IP knockdown did not result in a loss of ENO2 or gain of AR and FOXA1 expression in either PC-3 or DU145 cells (Supplemental Fig. 6C, D, E, and F). Thus, SLX4IP’s regulation of ALT hallmarks functions independent of NED, and by extension, AR-independent CRPC reprogramming.

## DISCUSSION

Presently, there exists a lack of targeted therapeutic options for patients with AR-independent CRPC, which includes therapeutically-acquired neuroendocrine prostate cancer^6^ and double-negative prostate cancer.^69^ Here, we report that SLX4IP-dependent ALT hallmarks are necessary for survival of AR-independent CRPC cell lines *in vitro* and *in vivo*. As our observations have been made in prostate cancer cell lines and correlated with gene expression studies in human tissues, whether ALT is the predominant TMM in AR-independent CRPC and whether ALT emerges as a consequence of androgen deprivation remains to be evaluated in patient tissues. Similarly, the independence of ALT and NED should be further confirmed in additional models of prostate cancer progression. Several *in vitro* models undergo AR-independent reprogramming following acquisition of resistance to second-generation antiandrogens indicated for CRPC treatment.^70^ Investigation of SLX4IP-mediated ALT hallmarks in these models of therapy-acquired NED would elucidate if this independence is consistent across AR-independent CRPC. These investigations would complement our findings that SLX4IP driven ALT-mediated telomere maintenance functions independent of AR loss and NED despite the reliance of AR-independent CRPC on this TMM.

The inability of ALT-dependent AR-independent CRPC to overcome SLX4IP loss, and subsequent induction of senescence, provides an elegant therapeutic target, but the knowledge void around ALT’s molecular mechanism must first be addressed. The ALT mechanism’s inherent complexity makes detailed pathway characterization challenging, thus the exact protein network required remains elusive. ALT is an HR-dependent replication pathway that requires the hijacking of already present DNA repair mechanisms to the telomere.^23^ This is unsurprising with lack of functional p53 and abundance of telomere dysfunction induced foci being a common ALT-positive occurrence.^23,71^ Additionally, insufficient telomere capping by shelterin protein TRF2 is permissive for telomeric recognition by ALT-associated DNA damage proteins.^72^ Collectively, these data suggest any protein involved in telomere homeostasis or DNA repair and replication may be involved in ALT activity. Interestingly, shelterin components TRF1, TRF2, RAP1 and TIN2 are required for APB formation despite the need for deficient telomere capping for ALT to occur.^24,72^ Therefore, a fine balance of telomere capping and shelterin sheathing is necessary to promote telomeric recombination while allowing recruitment of ALT mediators. One essential TRF2-mediated interaction is the SLX4 scaffold harboring nucleases responsible for the resolution of telomeric intermediate structures generated during HR.^35,36^ SLX4 is of particular interest as it binds SLX4IP, which we have described in our studies as a key regulator of ALT hallmarks in AR-independent CRPC.^36^ Together these previously identified regulators and many others contribute to ALT-mediated telomere elongation but these molecular players are also essential for genomic stability in non-malignant cellular populations. Therefore, identification of novel ALT regulators unique to malignant telomere maintenance is necessary to define therapeutically advantageous targets for AR-independent CRPC.

Our studies describe SLX4IP as a necessary regulator of ALT, that when removed induces telomere shortening and senescence separate of AR-independent reprogramming. Despite our studies and others describing the importance of SLX4IP in TMM shifts,^38,39^ the only known function of SLX4IP is through interactions with SLX4.^36^ As a presumed scaffold protein, it is possible SLX4IP assists SLX4 and other ALT mediators to localize at the telomere. As such, the identification of undefined interacting partners and post-translational regulators of SLX4IP may also prove therapeutically valuable. Potentially acting upon SLX4IP, the SLX4 nuclease complex functions as a SUMO E3 ligase in addition to its function in HR^73^, and perhaps analysis of post-translational modifications regulating the function or degradation of SLX4IP may reveal targets upstream of SLX4IP-mediated ALT hallmarks that can be exploited therapeutically.

Lastly, the dichotomous regulation of ALT and telomerase in CRPC described in our studies warrants further investigation. Several groups have demonstrated androgens transcriptionally regulate *TERT* in primary prostate cancer.^29,74,75^ Following progression to CRPC, AR mutants and splice variants are capable of differentially regulating telomerase expression.^75^ Furthermore, these AR mutants and splice variants coexist *in vitro* and *in vivo* making telomerase expression the collective output of AR perturbations present.^4^ Despite this complexity, AR-mediated regulation of telomerase supports our data demonstrating the reliance of AR-independent CRPC on SLX4IP-mediated ALT hallmarks. It is possible that either AR suppresses *SLX4IP* transcriptionally or TERT, shown to have noncanonical extra-telomeric functions, interferes with SLX4IP.^76^ Following progression to AR-independent CRPC, we have shown SLX4IP is free to promote ALT hallmarks as a backup following loss of telomerase. Interestingly, the AR-independent DU145 cell line exhibited rebound telomerase expression and activity following ALT inhibition despite maintained AR loss. This compensatory effect was described in other malignant investigations as well,^39^ but why this varies between two AR-independent CRPC cell lines remains undetermined. Nonetheless, the observed induction of telomerase was incapable of filling the TMM deficiency in the timeframe studied and both cell lines underwent telomere shortening and senescence. Because of this attempted resistance to ALT hallmark loss, defining mechanisms regulating both SLX4IP and compensatory telomerase in AR-independent CRPC will be crucial for describing the purpose of TMM shifts associated with disease progression

In summary, AR-independent CRPC exhibits ALT hallmarks and limited telomerase involvement while CRPC retaining AR lacks ALT hallmarks and uses telomerase *in vitro*. ALT hallmarks correlate with elevated SLX4IP expression and introduction of exogenous SLX4IP induces ALT hallmarks and reduces telomerase activity. Additionally, reduction of SLX4IP in AR-independent CRPC renders a telomeric environment lacking ALT hallmarks triggering senescence. This phenotype was recapitulated *in vivo* resulting in reduced tumor volume. With androgen deprivation, AR-dependent CRPC loses telomerase expression and activity and gains ALT hallmarks in an SLX4IP-dependent manner. Moreover, SLX4IP-mediated ALT hallmarks and AR-independent regulatory networks function independently. Together, these data reveal SLX4IP-mediated ALT hallmarks as a unique mechanism necessary for the replicative immortality of difficult to treat AR-independent CRPC.

## ACKNOWLEDGEMENTS

The authors would like to thank all members of the Taylor and Grabowska labs for helpful comments and suggestions related to this investigation. The Grabowska lab is supported by start-up funds (MMG) provided by the Case Research Institute, a joint venture between University Hospitals and Case Western Reserve University. The Taylor Lab is supported by the NIH (R01 GM133841 and CA240993 to DJT). We would also like to acknowledge the Molecular Therapeutics Training Program (T32 GM008056 to TLM).

## AUTHOR CONTRIBUTIONS

TLM, MMG, and DJT designed research; TLM performed research; WNA assisted with *in vivo* study; TLM, MMG, and DJT analyzed data and wrote the manuscript.

