## Supplemental Data for "SLX4IP-mediated telomere maintenance is essential for androgen receptor-independent castration-resistant prostate cancer"

**SUPPLEMENTAL INFORMATION**

**REFERENCES CONT.**

**A.**

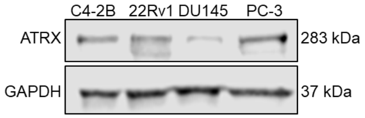


**B.**

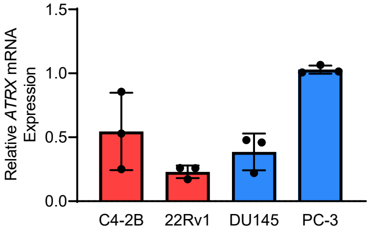


**Supplemental Figure 1: (A)** Relative ATRX protein expression across CRPC cell lines. **(B)** Relative *ATRX* mRNA expression across CRPC cell lines. Data represented as mean±SD; n=3; *p<0.05; **p<0.01; ***p<0.001; ****p<0.0001.

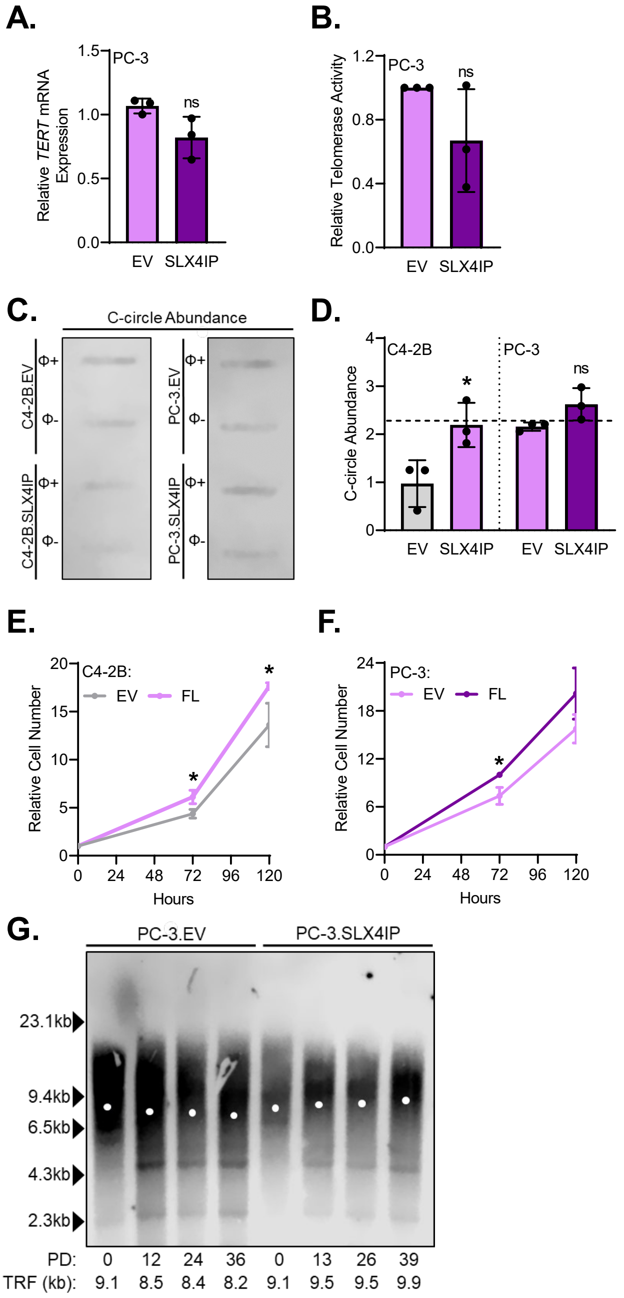


**Supplemental Figure 2:** **(A)** Relative *TERT* mRNA expression and **(B)** relative telomerase activity in PC-3 cells with stable SLX4IP overexpression. **(C)** Representative dot blot demonstrating the presence of telomeric C-circles in C4-2B and PC-3 cells with SLX4IP overexpression. Φ+ lane indicates reactions incubated with Φ29 polymerase for telomeric C-circle amplification. Φ- lane indicates control reaction lacking polymerase to account for background telomeric signal. **(D)** Quantification of C-circle abundance defined as the signal ratio of Φ+ reaction to Φ- reaction. Dotted line at y=2 indicates threshold signal ratio suggesting ALT activity. **(E)** Using trypan blue exclusion, relative cell number was followed over 120 hours for C4-2B and **(F)** PC-3 cells with SLX4IP overexpression. **(G)** Telomere restriction fragment analysis demonstrating telomere length changes over 45 days in PC-3 cells overexpressing SLX4IP versus control. PD: Calculated population doublings. Data represented as mean±SD; n=3; $*p<0.05$; $**p<0.01$; $***p<0.001$; $****p<0.0001$.

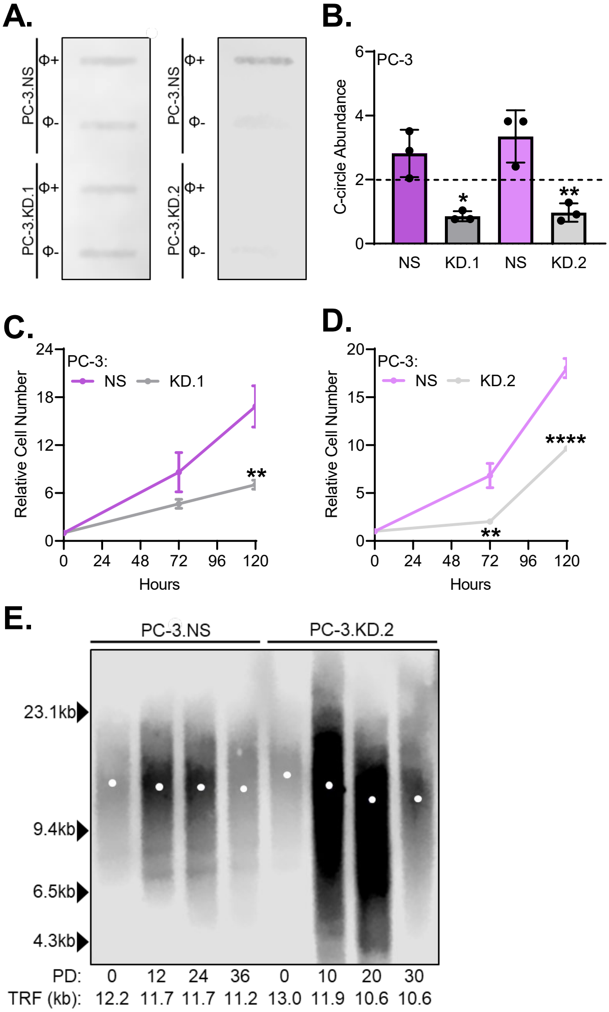


**Supplemental Figure 3: (A)** Representative dot blot demonstrating the presence of telomeric C-circles in PC-3 cells with SLX4IP knockdown. Φ+ lane indicates reactions incubated with Φ29 polymerase for telomeric C-circle amplification. Φ- lane indicates control reaction lacking polymerase to account for background telomeric signal. **(B)** Quantification of C-circle abundance defined as the signal ratio of Φ+ reaction to Φ- reaction. Dotted line at y=2 indicates threshold signal ratio suggesting ALT activity. **(C)** Using trypan blue exclusion, relative cell number was followed over 120 hours in PC-3.NS versus KD.1 or **(D)** KD.2 cells. **(E)** Telomere restriction fragment analysis demonstrating telomere length changes over 45 days in PC-3 cells with SLX4IP knockdown (KD.2) versus control. PD: Calculated population doublings. Data represented as mean+SD; n=3; *p<0.05; **p<0.01; ***p<0.001; ****p<0.0001.

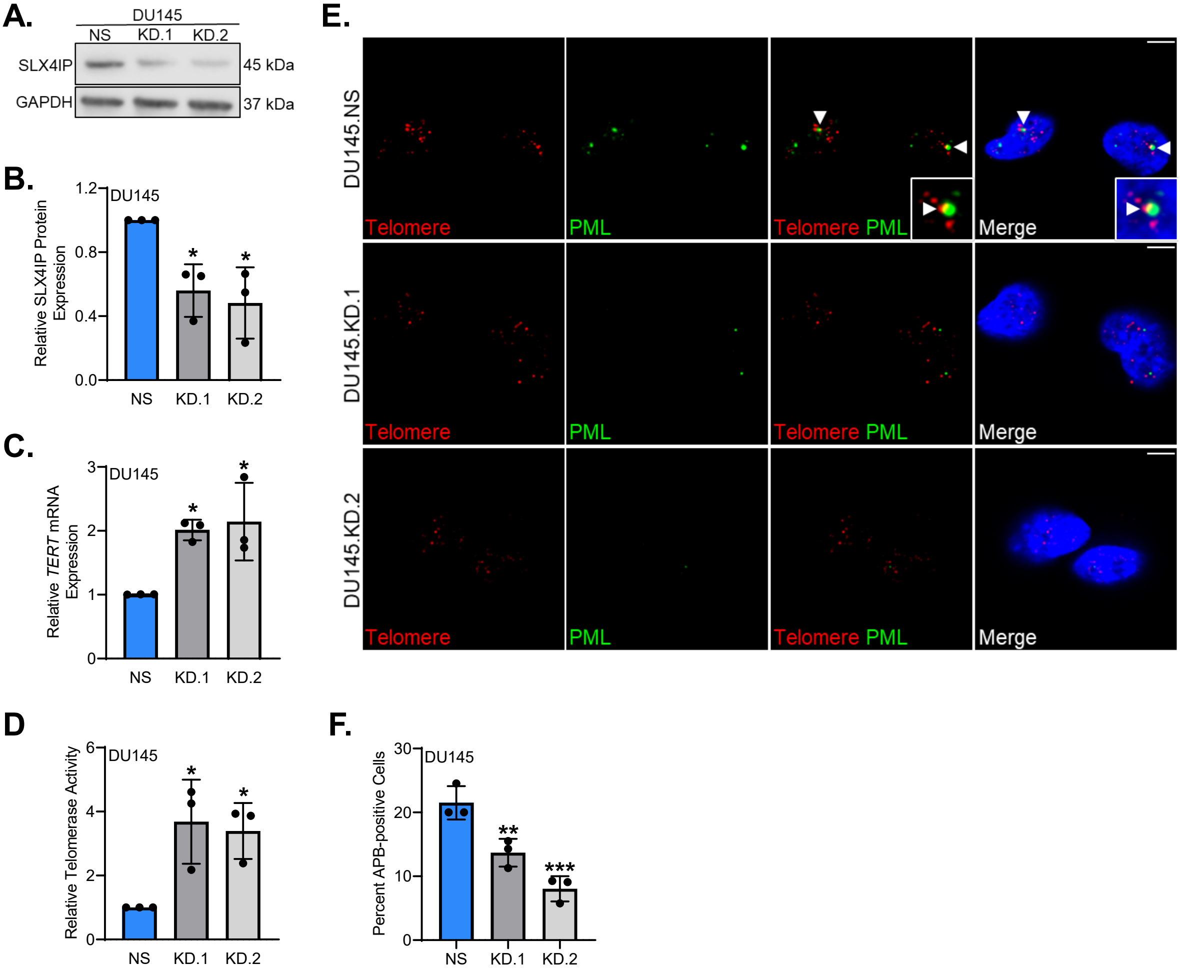


**Supplemental Figure 4: (A)** Confirmation of stable SLX4IP knockdown using two shRNAs (KD.1, KD.2) in DU145 cells with scrambled shRNA control (NS) at the protein level. **(B)** Quantification of (A). **(C)** Relative *TERT* mRNA expression and **(D)** relative telomerase activity following SLX4IP knockdown in DU145 cells. **(E)** Representative IF-FISH images demonstrating the presence or absence of APBs at telomeres (arrowheads) in DU145 cells with SLX4IP knockdown. Scale bar: 5 µm. **(F)** Quantification of IF-FISH images for percent APB-positive cells. Data represented as mean+SD; n=3; *p<0.05; **p<0.01; ***p<0.001; ****p<0.0001.

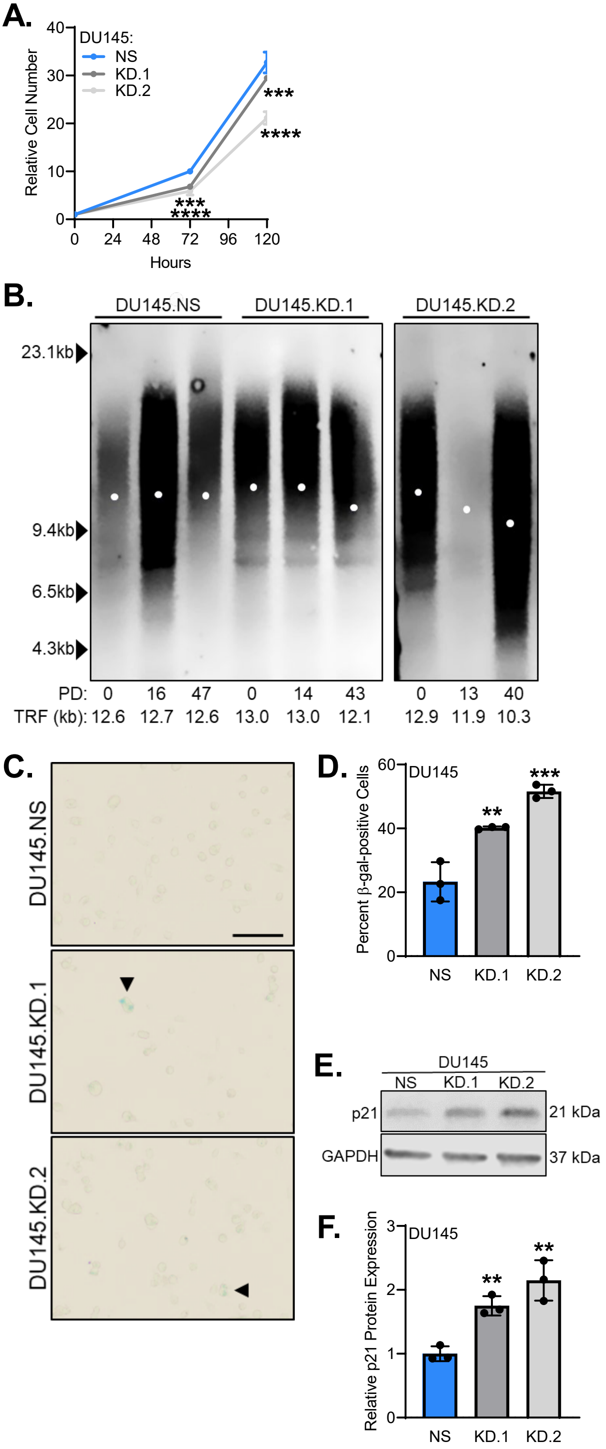


**Supplemental Figure 5: (A)** Using trypan blue exclusion, relative cell number was followed over 120 hours for DU145 cells with SLX4IP knockdown (KD.1, KD.2). **(B)** Telomere restriction fragment analysis demonstrating telomere length changes over 45 days in DU145 cells with SLX4IP knockdown (KD.1, KD.2) versus control. PD: Calculated population doublings. **(C)** Representative bright field images demonstrating β-galactosidase staining (arrowheads) for senescence in DU145 cells with stable knockdown of SLX4IP. Scale bar: 20 μm. **(D)** Quantification of (C). **(E)** Relative p21 expression following knockdown of SLX4IP in DU145 cells. **(F)** Quantification of (E). Data represented as mean±SD; n=3; *$p<0.05$; **$p<0.01$; ***$p<0.001$; ****$p<0.0001$.

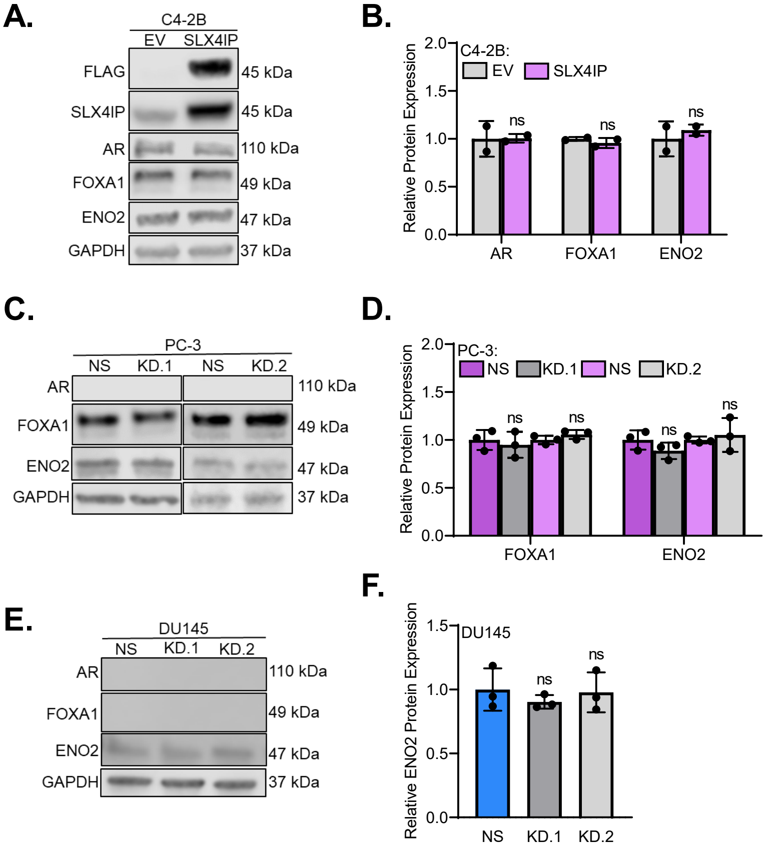


**Supplemental Figure 6: (A)** Changes in AR, FOXA1, and ENO2 protein expression in C4-2B transiently overexpressing SLX4IP. n=2 **(B)** Quantification of (A). **(C)** Changes in AR, FOXA1 and ENO2 protein expression in PC-3 cells with stable knockdown of SLX4IP. **(D)** Quantification of (C). **(E)** Changes in AR, FOXA1 and ENO2 protein expression in DU145 cells with stable knockdown of SLX4IP. **(F)** Quantification of (E) Data represented as mean±SD; n=3; *p<0.05; **p<0.01; ***p<0.001; ****p<0.0001.
